# Single-cell whole-brain imaging and network analysis provide evidence of the three-stage hypothesis of addiction

**DOI:** 10.1101/471847

**Authors:** Adam Kimbrough, Daniel J. Lurie, Andres Collazo, Max Kreifeldt, Harpreet Sidhu, Mark D’Esposito, Candice Contet, Olivier George

## Abstract

Three main theories of the neurobiology of addiction have been proposed: (1) incentive salience mediated by a brainstem-striatal network, (2) habit mediated by a cortico-striato-thalamic network, and (3) hedonic allostasis mediated by an extended amygdala network. Efforts have been made to reconcile these theories within a three-stage model, but the relevance of each theory remains controversial. We tested the validity of each theory with a single dataset using unbiased single-cell whole-brain imaging and data-driven analyses of neuronal activity in a mouse model of alcohol use disorder. Abstinence in alcohol dependent mice decreased brain modularity and resulted in clustering of brain regions that correspond to each stage of the three-stage theory of addiction. Furthermore, we identified several brain regions whose activity highly predicted addiction-like behaviors and “hub” regions that may drive neural activation during abstinence. These results validate the three-stage theory of addiction and identify potential target regions for future study.

## Introduction

Three major theories have been proposed to explain the neurobiological basis of addiction and substance abuse (**Fig. 1**): (1) the incentive salience theory of craving and wanting drug, which is thought to involve the mesocorticolimbic dopamine system (Robinson and Berridge, 2008, Berridge and Robinson, 2016, Robinson and Berridge, 1993), (2) the habit/compulsivity theory of Pavlovian conditioning and the formation of habitual drug use, which is thought to involve the prefrontal cortex (PFC) and striatum (Everitt and Robbins, 2005, Everitt, 2014), and (3) the hedonic allostasis theory of negative reinforcement that drives drug intake and escalation, which is thought to involve the extended amygdala (Koob, 2008c, George et al., 2012a). Efforts have been made to reconcile these theories using a three-stage theory of “craving, binge, and abstinence,” which is thought to involve a combination of many of the neural systems that are encompassed in the other theories (Koob and Volkow, 2016, Koob and Le Moal, 1997, Koob and Volkow, 2010). However, this topic has remained controversial (George et al., 2014, Badiani, 2014, Piazza and Deroche-Gamonet, 2013, Piazza and Deroche-Gamonet, 2014, Wise and Koob, 2014), with several treaties suggesting that one or more theories are not relevant or not required for the development of addiction (Pascoli et al., 2015, Lovic et al., 2011). Determining the brain regions that are associated with addiction, the ways in which they are co-activated, and the ways in which they fit into the theoretical framework of addiction using an unbiased approach would be a major step forward in the field.

**Figure 1.**
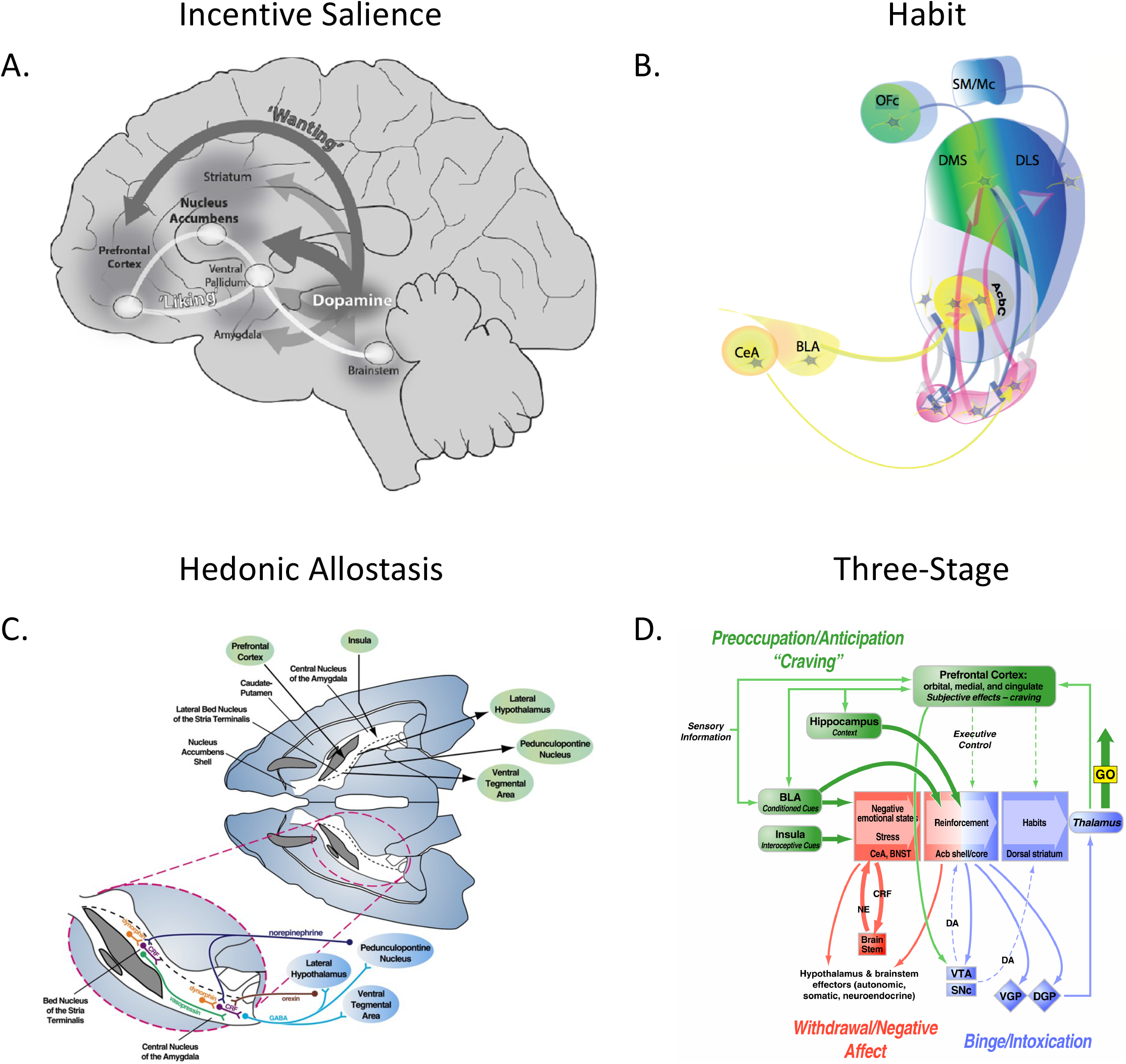
Theories of the neurobiology of addiction. **A**. Incentive salience theory of craving and wanting drug, which is thought to involve the mesocorticolimbic dopamine system **B**. Habit/compulsivity theory of Pavlovian conditioning and formation of habitual drug use, which is thought to involve corticostriatal loops **C**. Hedonic allostasis theory producing symptoms of negative affect, which is thought to involve the striatum and extended amygdala. **D**. The three-stage theory of “craving, binge, and abstinence,” which is thought to involve a combination of many of the neural systems that are encompassed in the other theories. Images reproduced with permission from (Koob et al., 2008, Koob, 2008d, Berridge and Robinson, 2016, Everitt, 2014).

Novel whole-brain imaging approaches, such as CLARITY, immunolabeling-enabled 3D imaging of solvent cleared organs (iDISCO), ultimate DISCO (uDISCO), and others (Chung and Deisseroth, 2013, Renier et al., 2014, Azaripour et al., 2016, Seo et al., 2016, Renier et al., 2016), have provided opportunities to expand our knowledge of functional neural circuitry in animal models of addiction in an unbiased manner. Using single-cell whole-brain imaging, data-driven hierarchical clustering, and graph theory analyses, we can identify specific modules (i.e., discrete clusters of brain regions) at single-cell resolution that show patterns of co-activation during alcohol abstinence. By clustering brain regions into groups that share co-activation profiles, we can determine the ways in which the organization of the brain fits into theoretical frameworks of addiction. Previous attempts have been made to identify clusters of brain regions that are associated with alcohol abstinence through literature review (George and Koob, 2010, Noori et al., 2012), but still unclear is whether the posited clusters represent an accurate picture of the full set of brain regions that are associated with alcohol abstinence.

Thus, we sought to use a mouse model of alcohol dependence to determine which, if any, theories of the neurobiology of addiction fit best with the neuronal networks that are associated with abstinence from alcohol dependence. To accomplish this, we (1) assessed the predictive value of neuronal activity from individual brain regions to key behavioral outcomes that are associated with dependence (alcohol drinking, irritability-like behavior, and digging behavior), (2) determined whether the network of co-activated brain regions is altered by alcohol abstinence by identifying clusters that are specifically associated with alcohol abstinence, and (3) determined potentially key or “hub” brain regions (e.g., regions that present high levels of inter-or intracluster connectivity) that are associated with alcohol abstinence using graph theory.

## Results

### Alcohol-dependent mice exhibit increases in alcohol drinking, irritability-like behavior, and digging behavior

We used a well-validated mouse model of alcohol dependence, the two-bottle choice (2BC)/chronic intermittent ethanol (CIE) vapor paradigm (Contet et al., 2011, Kreifeldt et al., 2013, Becker and Lopez, 2004, Gorini et al., 2013), to explore the relationship between behavioral and neural effects of alcohol abstinence. We assessed alcohol intake in alcohol-dependent (2BC/CIE) and nondependent (2BC/Air) mice and assessed irritability-like behavior and digging behavior in 2BC/CIE, 2BC/Air, and alcohol-naive mice. We also identified functional co-activation networks during alcohol abstinence in alcohol-dependent mice and compared them to networks from alcohol-nondependent and naive mice. For an experimental outline, see **Fig. 2**.

**Figure 2.**
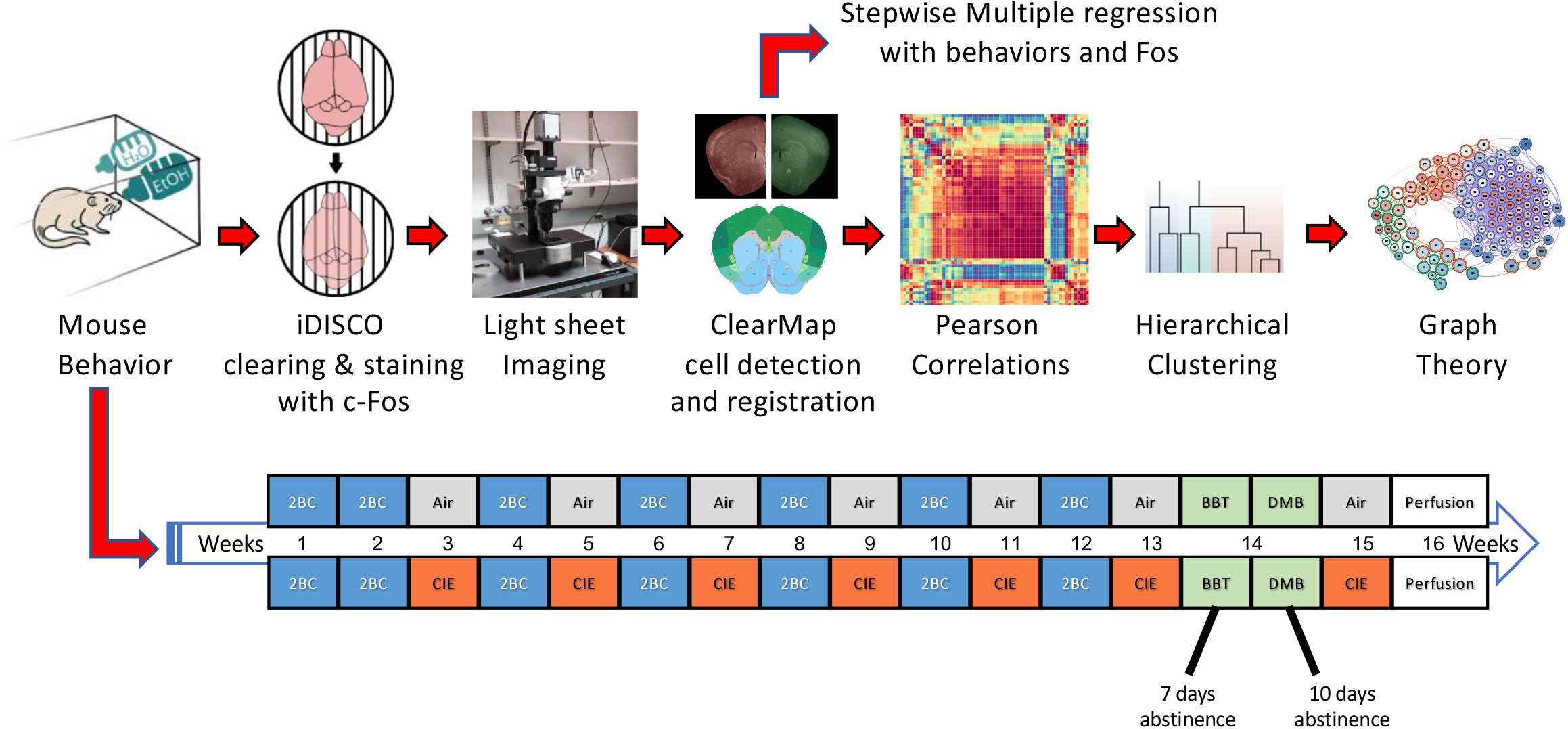
Experimental design. The mice underwent the 2BC/CIE paradigm. The 2BC/CIE paradigm involves alternating weeks of 2BC (highlighted in blue) and CIE/Air (highlighted in red and gray, respectively). The mice received 2 weeks of baseline 2BC testing followed by five rounds of alternating CIE or Air weeks and 2BC weeks. The mice then received a 6^th^ week of CIE/Air and were then tested for irritability-like and digging behaviors 7 and 10 days, respectively, into abstinence (highlighted in green). The mice then received an additional week of CIE/Air, and brains were collected 7 days after the last vapor exposure with no intervening behavioral testing. Brains were collected during the time of day that the mice would normally undergo a 2BC session. Brains were then immunostained for Fos and cleared using the iDISCO procedure. Brains were then imaged and analyzed further to identify brain regions that potentially contribute to behaviors and the functional networks for each treatment. Key brain regions in the alcohol abstinence network were then identified using graph theory. 2BC, two-bottle choice; CIE, chronic intermittent ethanol vapor; BBT, bottle-brush test of irritability-like behavior; DM, digging and marble burying.

### Alcohol dependence escalates voluntary alcohol consumption

After 2 weeks of baseline alcohol drinking and five rounds of alternating weeks of 2BC/CIE exposure, alcohol-dependent mice exhibited an increase in drinking compared with nondependent mice that had similar access to drinking but no exposure to alcohol vapor. The repeated-measures analysis of variance (ANOVA), with group (2BC/CIE and 2BC/Air) as the between-subjects factor and week (baseline and post-vapor intake) as the within-subjects factor, revealed a significant week × group interaction (*FF*_1,7_ = 13.9,*p* < 0.05) and a significant effect of group (*F*_1,7_ = 10.0, p < 0.05). The Student-Newman Keuls (SNK) *post hoc* test revealed that 2BC/CIE mice significantly escalated alcohol intake during post-vapor week 5 *vs*. their own baseline (2.8 ± 0.2 g/kg/2 h at baseline *vs*. 3.7 ± 0.3 g/kg/2 h at post-vapor week 5), and 2BC/CIE mice had significantly higher alcohol intake compared with 2BC/Air mice at postvapor week 5 (3.7 ± 0.3 g/kg/2 h for 2BC/CIE *vs*. 2.3 ± 0.2 g/kg/2 h for 2BC/Air at post-vapor week 5; **Fig. 3A**, see **Supplemental Fig. S1** for the complete time course of escalation).

**Figure 3.**
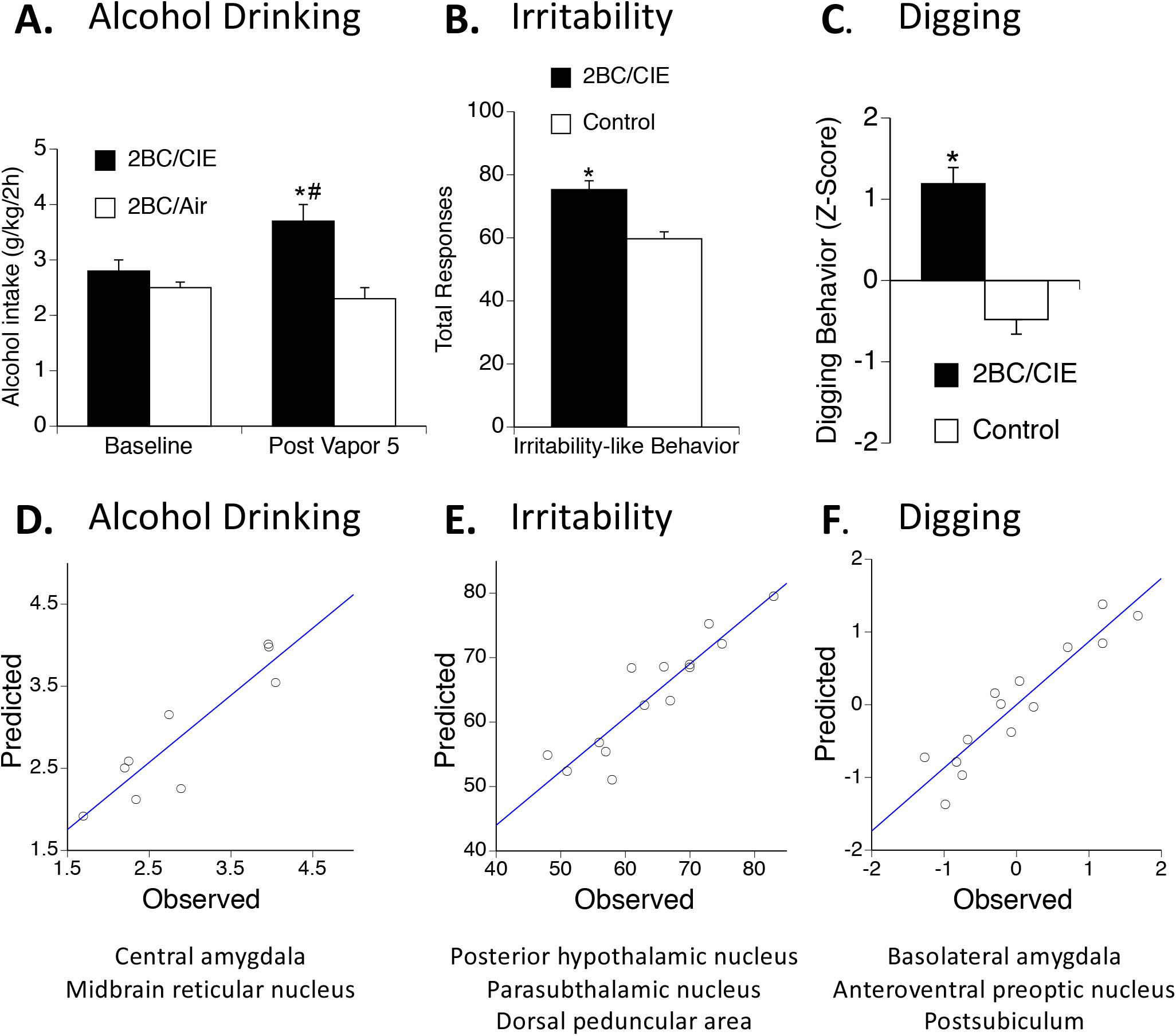
Alcohol drinking and abstinence behavior in 2BC/CIE *vs*. control mice and predicted *vs*. observed values for stepwise multiple regression analysis. **A**. 2BC/CIE mice (black bars) exhibited a significant increase in alcohol drinking during post-vapor week 5 2BC testing compared with alcohol drinking in 2BC/Air mice (white bars) and their own baseline intake. **B**. Irritability-like behavior. 2BC/CIE mice (black bar) exhibited a significant increase in total irritable-like responses compared with control mice (2BC/Air and naive; white bar). **C**. Digging behavior. 2BC/CIE mice (black bar) exhibited a significant increase in composite digging behavior (Z-score of digging behaviors) compared with control mice (2BC/Air and naive; white bar). \**p* < 0.05 (two-tailed), 2BC/CIE *vs*. 2BC/Air for alcohol drinking or control for irritabilitylike and digging behavior; ^#^*p* < 0.05 (two-tailed), 2BC/CIE post-vapor week 5 *vs*. 2BC/CIE baseline. **D-F**. Observed values for each behavior and predicted values for each behavior based on stepwise multiple regression analysis. The brain regions that were identified by the analysis as contributing to the behavior are listed below each graph.

### Irritability-like behavior

To evaluate affective dysfunction during alcohol abstinence, we measured irritability-like behavior. Irritability is a key symptom of alcohol abstinence in rodents (Somkuwar et al., 2017, Kimbrough et al., 2017, Sidhu et al., 2018) and a central feature of alcohol dependence in humans together with greater aggression and frustration (Miczek et al., 2015, Cardoso et al., 2006, Winward et al., 2014, Baars et al., 2013, Lubman et al., 1983). 2BC/Air mice and naive mice did not differ from each other in irritability-like behavior (60.8 ± 3.4 for 2BC/Air *vs*. 58.6 ± 3.2 for naive; *t* = 0.47, *p* > 0.05); therefore, we combined 2BC/Air and naive mice into a single control group. We found a significant increase in irritability-like behavior 1 week into abstinence in 2BC/CIE mice compared with control mice (75.3 ± 2.8 for 2BC/CIE *vs*. 59.7 ± 2.2 for control; *t* = 3.92, *p* < 0.005; **Fig. 3B**).

### Digging behavior

To further evaluate affective dysfunction during abstinence, we measured digging and marble-burying behavior (Sidhu et al., 2018, Jury et al., 2017, Pleil et al., 2015a, Rose et al., 2016, Umathe et al., 2008). Similar to irritability-like behavior, 2BC/Air and naive mice did not significantly differ from each other in their composite digging behavior (−0.70 ± 0.25 for 2BC/Air *vs*. −0.26 ± 0.14 for naive; *t* = 1.5, *p* > 0.05); therefore, we combined 2BC/Air and naive mice into a single control group. 2BC/CIE mice exhibited a significant increase in digging behavior 1 week into abstinence compared with control mice (1.19 ± 0.20 for 2BC/CIE *vs*. −0.48 ± 0.18 for control; *t* = 6.03, *p* < 0.0005; **Fig. 3C;** see **Supplemental Fig. S2** for individual digging behaviors).

### Neural activation predicts behaviors associated with alcohol abstinence

To evaluate neuronal recruitment during alcohol abstinence, we used single-cell whole-brain imaging using iDISCO with immunohistochemistry to detect immediate early gene c-*fos* expression as a proxy for neuronal activation (Renier et al., 2016, Renier et al., 2014). c-*fos* was chosen over other immediate early genes because its low baseline levels are optimal for detecting increases in neuronal activity and based on previous success using Fos with the iDISCO technique by Renier et al. (2014 and 2016).

### A subset of brain regions highly predicted alcohol abstinence-associated behaviors

To identify brain regions that are the most predictive of behaviors that are associated with alcohol abstinence, we performed stepwise multiple regression analysis using Fos counts from all brain regions relative to each behavioral measure (drinking, irritability-like behavior, and digging behavior). When alcohol drinking was analyzed, two brain regions (central nucleus of the amygdala [CeA] and midbrain reticular nucleus [MRN]) were identified as responsible for the majority of the variability (adjusted *R*^2^ = 0.757; *F*_2,6_ = 13.5, *p* < 0.05). The intercept *b* value was −2.04 ± 1.20. The CeA had a *b* value of 4.46 ± 0.87 (*p* < 0.05), and the MRN had a *b* value of −1.58 ± 0.60 (*p* < 0.05) for a formula of y = −2.04 + 4.46 * CEA – 1.58 * MRN. A plot of observed *vs*. predicted values for drinking based on Fos counts from the CeA and MRN is presented in **Fig. 3D**.

When irritability-like behavior was analyzed, three brain regions (posterior hypothalamic nucleus [PH], parasubthalamic nucleus [PSTN], and dorsal peduncular area [DP]) were identified as responsible for the majority of the variability (adjusted *R*^2^ = 0.785; *F*_3,10_ = 16.8, *p* < 0.005). The intercept had a *b* value of 37.75 ± 4.43. The PH had a *b* value of 8.65 ± 2.08 (*p* < 0.005), the PSTN had a *b* value of 11.93 ± 2.88 (*p* < 0.005), and the DP had a *b* value of 7.30 ± 2.81 (*p* < 0.05) for a formula of y = 37.75 + 8.65 * PH + 11.93*PSTN + 7.30*DP. A plot of observed *vs*. predicted values for irritability-like behavior based on Fos counts from the PH, PSTN, and DP is presented in **Fig. 3E**.

When composite digging behavior was analyzed, three brain regions (basolateral amygdala [BLA], anteroventral preoptic nucleus [AVP], and postsubiculum [POST]) were identified as responsible for the majority of the variability (adjusted *R*^2^ = 0.829; *F*_3,10_ = 22.0, *p* < 0.005). The intercept had a *b* value of −4.41 ± 0.56. The BLA had a *b* value of 1.47 ± 0.23 (*p* < 0.005), the AVP had a *b* value of 0.79 ± 0.22 (*p* < 0.005), and the POST had a *b* value of 0.54 ± 0.20 (*p* < 0.05) for a formula of y = −4.41 + 1.47*BLA + 0.79*AVP + 0.54*POST. A plot of observed *vs*. predicted values for composite digging behavior based on Fos counts from the BLA, AVP, and POST is presented in **Fig. 3F**. No observed values were found to be outliers in any of the behaviors based on Cook’s *D*.

### Identification of large-scale patterns of co-activation associated with alcohol abstinence Co-activation patterns are significantly altered by alcohol abstinence

Drug dependence may be driven by specific clusters/modules of brain regions, the coordinated activity of multiple modules throughout the brain, or some combination of the two (George and Koob, 2010, Noori et al., 2012). To explore whole-brain patterns of neural coactivation that is associated with each of the experimental conditions, we first visualized interregional Fos correlations for each condition. Correlation matrices were organized according to traditional anatomical groups from the Allen Mouse Brain Atlas (**Fig 4;** see **Table 1** and methods for the order in which brain regions are listed). When visualized this way, clear differences can be observed in co-activation patterns between mice in the 2BC/CIE and the control conditions (2BC/Air and naive; **Fig. 4A-C**). Overall, 2BC/CIE mice exhibited a higher level of cross-correlation between brain regions compared with the control conditions. However, one cluster of brain regions that included the lateral amygdala (LA), endopiriform nucleus (EP), BLA, CeA, and intercalated amygdala (IA) clearly stood out by being anticorrelated with most of the other brain regions (framed in **Fig. 4A-C**). To further identify the way in which this amygdala cluster is functionally connected to the rest of the brain, we compared correlation patterns of the amygdala cluster with the brain regions that exhibited the highest levels of correlation/anticorrelation by calculating the average correlation across each brain region group (**Fig. 5A, B**). Significant effect of group were found in all comparisons of the amygdala cluster to the other clusters that were examined: amygdala cluster intra-cluster correlation (excluding self-correlations; *F*_2,12_ = 93.8, *p* < 0.005), cortical amygdala/retrohippocampal cluster (cortical amygdalar nucleus posterior part [COAp], ENTm, entorhinal area lateral part [ENTl], and parasubiculum; *F*_2,12_ = 13.5, *p* < 0.005), PSTN/tuberal nucleus (TU; *F*_2,12_ = 14.0, *p* < 0.005), cortical cluster (orbitofrontal cortex [OFC], prefrontal cortex [PFC], and sensory/somatosensory cortex; *F*_2,12_ = 34.1, *p* < 0.005), hippocampal cluster (HIPP; field CA1, CA2, CA3, and dentate gyrus; *F*_2,12_ = 23.2, *p* < 0.005), thalamic cluster (THAL; major thalamic nuclei; *F*_2,12_ = 8.6, *p* < 0.005), hypothalamic cluster (HYPO; major hypothalamic nuclei; *F*_2,12_ = 35.0, *p* < 0.005), and interpeduncular nucleus (IPN) and ventral tegmental area (VTA; *F*_2,12_ = 4.0, *p* < 0.05; **Fig. 5B**).

**Figure 4.**
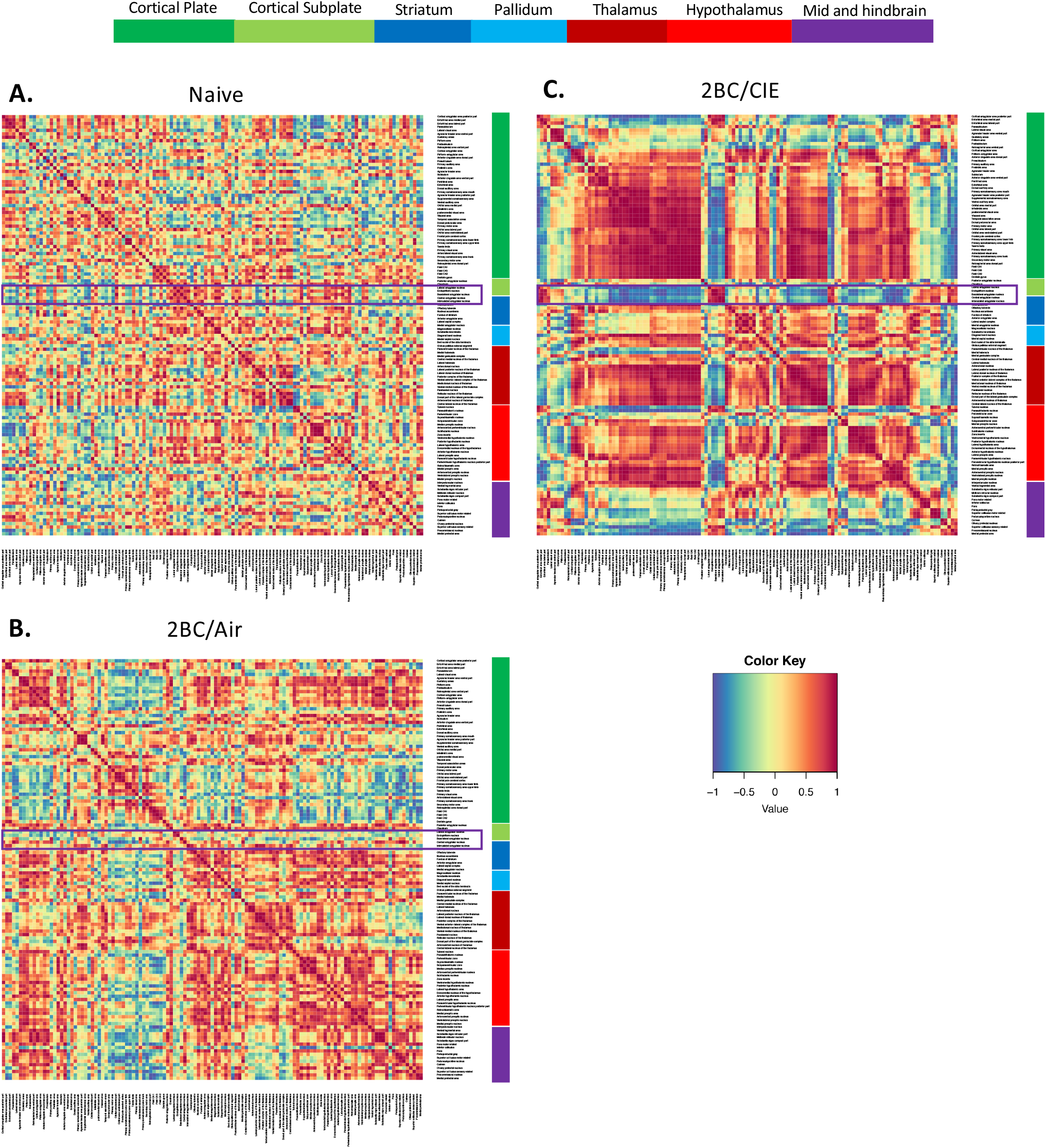
Interbrain regional Pearson correlation heatmaps for each treatment organized anatomically based on the Allen Mouse Brain Atlas. **A**. Correlation heatmap for naive mice. **B**. Correlation heatmap for 2BC/Air mice. **C**. Correlation heatmap for 2BC/CIE mice. Each heatmap is organized into color-coded anatomical groups: dark green (cortical plate), light green (cortical subplate), dark blue (striatum), light blue (pallidum), dark red (thalamus), light red (hypothalamus), purple (midbrain, hindbrain, and cerebellum). The region that is highlighted in purple on each heatmap represents an amygdala cluster of the heatmap shown in greater detail in Fig. 5.

**Figure 5.**
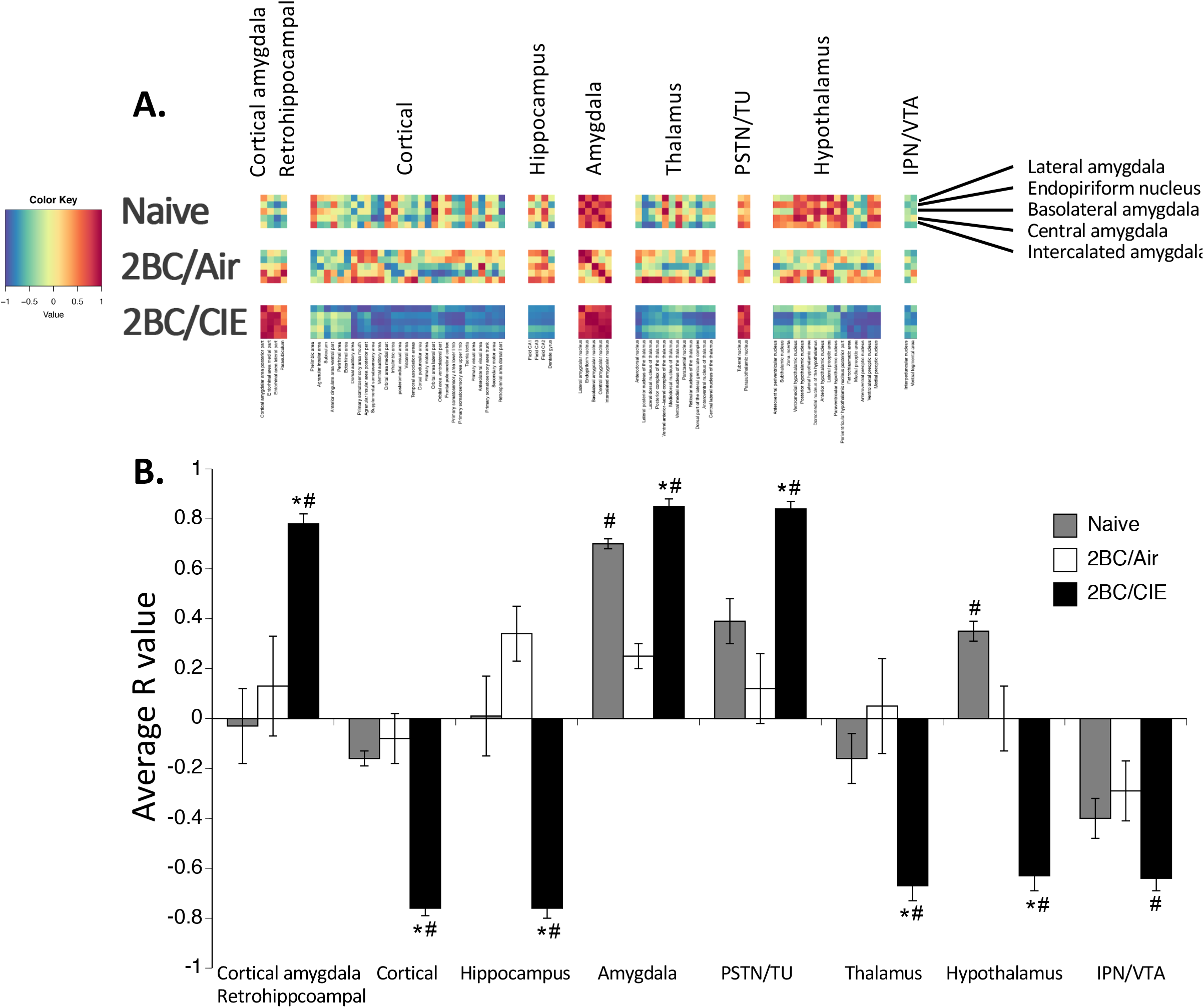
Comparison of Pearson correlations of an amygdala cluster *vs*. other regions. **A**. Cut out of correlations from Fig. 4 of brain regions compared with the amygdala cluster for each treatment. Individual region names are displayed at the bottom, and group names are displayed above each cluster. **B**. Average *R* values for 2BC/CIE (black bars), 2BC/Air (white bars), and naive (gray bars) for each cluster *vs*. the amygdala cluster. 2BC/CIE mice showed either a strong positive or strong negative correlation with each cluster. Naive mice only showed a strong positive correlation with the amygdala cluster. 2BC/Air mice showed no strong positive or negative correlation with any cluster. \**p* < 0.05, *vs*. naive; ^#^*p* < 0.05, *vs*. 2BC/Air.

**Table 1.** Brain region names, abbreviations, Allen Brain atlas grouping, abstinence cluster, participation coefficient, and within-module degree Z score values for abstinence network.

| Brain Region | Abbreviation | Allen Group Name | Abstinence Cluster | PC | WMDz |
| --- | --- | --- | --- | --- | --- |
| Agranular insular area | AI | Cortical Plate | C | 0 | -1.46 |
| Agranular insular area, posterior part | AIp | Cortical Plate | C | 0 | -0.51 |
| Agranular insular area, ventral part | AIv | Cortical Plate | B | 0.67 | -1.02 |
| Anterior cingulate area, dorsal part | ACA<sub>d</sub> | Cortical Plate | C | 0.26 | -0.77 |
| Anterior cingulate area, ventral part | ACA<sub>v</sub> | Cortical Plate | C | 0.25 | -1.83 |
| Anterolateral visual area | VIS<sub>al</sub> | Cortical Plate | C | 0 | 0.73 |
| Cortical amygdalar area | COA | Cortical Plate | B | 0.49 | 0.74 |
| Cortical amygdalar area, posterior part | COA<sub>p</sub> | Cortical Plate | A | 0.16 | 1.61 |
| Dentate gyrus | DG | Cortical Plate | C | 0 | 1.13 |
| Dorsal auditory area | AUD<sub>d</sub> | Cortical Plate | C | 0 | -0.37 |
| Dorsal peduncular area | DP | Cortical Plate | C | 0 | 0.58 |
| Ectorhinal area | ECT | Cortical Plate | C | 0.08 | -0.12 |
| Entorhinal area, lateral part | ENT<sub>l</sub> | Cortical Plate | A | 0.44 | -0.12 |
| Entorhinal area, medial part | ENT<sub>m</sub> | Cortical Plate | A | 0.45 | 0.41 |
| Field CA1 | CA1 | Cortical Plate | C | 0 | 1.33 |
| Field CA2 | CA2 | Cortical Plate | C | 0 | 1.41 |
| Field CA3 | CA3 | Cortical Plate | C | 0 | 1.39 |
| Frontal pole, cerebral cortex | FRP | Cortical Plate | C | 0 | 0.96 |
| Gustatory areas | GU | Cortical Plate | A | 0.49 | -1.26 |
| Infralimbic area | ILA | Cortical Plate | C | 0 | 0.35 |
| Lateral visual area | VIS<sub>l</sub> | Cortical Plate | B | 0.65 | -1.04 |
| Orbital area, lateral part | ORB<sub>l</sub> | Cortical Plate | C | 0 | 1.13 |
| Orbital area, medial part | ORB<sub>m</sub> | Cortical Plate | C | 0 | -0.03 |
| Orbital area, ventrolateral part | ORB<sub>vl</sub> | Cortical Plate | C | 0 | 1.10 |
| Parasubiculum | PAR | Cortical Plate | B | 0.27 | 0.52 |
| Perirhinal area | PERI | Cortical Plate | C | 0.06 | -0.85 |
| Piriform area | PIR | Cortical Plate | A | 0.50 | -0.88 |
| Piriform-amygdalar area | PAA | Cortical Plate | C | 0.30 | -0.60 |
| Posteromedial visual area | VIS<sub>pm</sub> | Cortical Plate | C | 0 | 0.21 |
| Postsubiculum | POST | Cortical Plate | B | 0.64 | -1.55 |
| Prelimbic area | PL | Cortical Plate | C | 0 | -0.32 |
| Presubiculum | PRE | Cortical Plate | C | 0.31 | -1.31 |
| Primary auditory area | AUD<sub>p</sub> | Cortical Plate | C | 0.44 | -2.18 |
| Primary motor area | Mop | Cortical Plate | C | 0 | 0.57 |
| Primary somatosensory area, lower limb | SSp-ll | Cortical Plate | C | 0 | 0.93 |
| Primary somatosensory area, mouth | SSp-m | Cortical Plate | C | 0 | -0.15 |
| Primary somatosensory area, trunk | SSp-tr | Cortical Plate | C | 0 | 1.09 |
| Primary somatosensory area, upper limb | SSp-ul | Cortical Plate | C | 0 | 0/86 |
| Primary visual area | VISp | Cortical Plate | C | 0 | 0.73 |
| Retrosplenial area, dorsal part | RSPd | Cortical Plate | C | 0 | 1.06 |
| Retrosplenial area, ventral part | RSPv | Cortical Plate | A | 0.50 | -1.66 |
| Secondary motor area | MOs | Cortical Plate | C | 0 | 1.11 |
| Subiculum | SUB | Cortical Plate | C | 0.13 | -1.75 |
| Supplemental somatosensory area | SSs | Cortical Plate | C | 0 | 0.68 |
| Taenia tecta | TT | Cortical Plate | C | 0 | 0.89 |
| Temporal association areas | Tea | Cortical Plate | C | 0 | 0.51 |
| Ventral auditory area | AUDv | Cortical Plate | C | 0 | 0.46 |
| Visceral area | VISC | Cortical Plate | C | 0 | 0.61 |
| Basolateral amygdalar nucleus | BLA | Cortical Subplate | A | 0 | 1.25 |
| Clastrum | CLA | Cortical Subplate | A | 0.47 | -1.28 |
| Endopiriform nucleus | EP | Cortical Subplate | A | 0.17 | 0.79 |
| Lateral amygdalar nucleus | LA | Cortical Subplate | A | 0.32 | 0.36 |
| Posterior amygdalar nucleus | PA | Cortical Subplate | C | 0.42 | -2.43 |
| Anterior amygdalar area | AAA | Striatum | B | 0.48 | -0.04 |
| Caudoputamen | CP | Striatum | C | 0 | 1.26 |
| Central amygdalar nucleus | CEA | Striatum | A | 0.47 | 0.81 |
| Fundus of striatum | FS | Striatum | B | 0.12 | 1.29 |
| Intercalated amygdalar nucleus | IA | Striatum | A | 0.35 | 1.37 |
| Lateral septal complex | LSX | Striatum | B | 0.50 | 0.96 |
| Medial amygdalar nucleus | MEA | Striatum | B | 0.50 | 1.01 |
| Nucleus accumbens | ACB | Striatum | C | 0.38 | -1.85 |
| Olfactory tubercle | OT | Striatum | C | 0.07 | -0.99 |
| Bed nuclei of the stria terminalis | BST | Pallidum | C | 0.33 | -0.84 |
| Diagonal band nucleus | NDB | Pallidum | C | 0 | 0.34 |
| Globus pallidus external segment | GPe | Pallidum | C | 0.21 | -0.66 |
| Magnocellular nucleus | MA | Pallidum | B | 0.41 | -0.60 |
| Medial septal nucleus | MS | Pallidum | C | 0 | -0.71 |
| Substantia innominata | SI | Pallidum | B | 0.47 | 1.26 |
| Anterodorsal nucleus | AD | Thalamus | C | 0 | 0.77 |
| Anteroventral nucleus of the thalamus | AV | Thalamus | C | 0 | 0.94 |
| Central lateral nucleus of the thalamus | CL | Thalamus | C | 0 | 0.92 |
| Central medial nucleus of the thalamus | CM | Thalamus | C | 0.32 | -0.60 |
| Lateral geniculate complex, dorsal part | LGd | Thalamus | C | 0 | 1.10 |
| Lateral dorsal nucleus of the thalamus | LD | Thalamus | C | 0 | 1.14 |
| Lateral habenula | LH | Thalamus | C | 0.22 | -1.68 |
| Lateral posterior nucleus of the thalamus | LP | Thalamus | C | 0 | 1.07 |
| Medial geniculate complex | MG | Thalamus | C | 0.32 | -1.83 |
| Medial habenula | MH | Thalamus | A | 0 | -1.33 |
| Mediodorsal nucleus of the thalamus | MD | Thalamus | C | 0.10 | 0.58 |
| Parataenial nucleus | PT | Thalamus | C | 0.04 | 0.34 |
| Paraventricular nucleus of the thalamus | PVT | Thalamus | B | 0.40 | -0.64 |
| Posterior complex of the thalamus | PO | Thalamus | C | 0 | 0.59 |
| Reticular nucleus of the thalamus | RT | Thalamus | C | 0.04 | 0.26 |
| Ventral anterior-lateral complex of the thalamus | VAL | Thalamus | C | 0 | 1.00 |
| Ventral medial nucleus of the thalamus | VM | Thalamus | C | 0.18 | 0.13 |
| Anterior hypothalamic nucleus | AHN | Hypothalamus | C | 0.24 | -0.22 |
| Anteroventral periventricular nucleus | AVPV | Hypothalamus | C | 0.08 | 0.05 |
| Anteroventral preoptic nucleus | AVP | Hypothalamus | C | 0 | -0.05 |
| Dorsomedial nucleus of the hypothalamus | DMH | Hypothalamus | C | 0.23 | -0.15 |
| Lateral hypothalamic area | LHA | Hypothalamus | C | 0.31 | -0.71 |
| Lateral preoptic area | LPO | Hypothalamus | C | 0 | -0.37 |
| Medial preoptic area | MPO | Hypothalamus | C | 0 | 0.75 |
| Medial preoptic nucleus | MPN | Hypothalamus | C | 0 | 0.23 |
| Median preoptic nucleus | MEPO | Hypothalamus | B | 0.32 | -1.82 |
| Parasubthalamic nucleus | PSTN | Hypothalamus | A | 0 | 1.12 |
| Paraventricular hypothalamic nucleus | PVH | Hypothalamus | C | 0 | -0.45 |
| Periventricular hypothalamic nucleus, posterior part | PVp | Hypothalamus | C | 0 | -1.41 |
| Periventricular zone | PVZ | Hypothalamus | B | 0.13 | 1.12 |
| Posterior hypothalamic nucleus | PH | Hypothalamus | C | 0.22 | -0.82 |
| Retrochiasmatic area | RCH | Hypothalamus | C | 0.11 | -1.56 |
| Subparaventricular zone | SBPV | Hypothalamus | C | 0.18 | -1.95 |
| Subthalamic nucleus | STN | Hypothalamus | C | 0.04 | 0.28 |
| Suprachiasmatic nucleus | SCH | Hypothalamus | B | 0.50 | -1.55 |
| Tuberal nucleus | TU | Hypothalamus | A | 0.49 | -0.31 |
| Ventrolateral preoptic nucleus | VLPO | Hypothalamus | C | 0 | 0.32 |
| Ventromedial hypothalamic nucleus | VMH | Hypothalamus | C | 0.31 | -0.92 |
| Zona incerta | ZI | Hypothalamus | C | 0.04 | 0.19 |
| Inferior colliculus | IC | Midbrain | B | 0.58 | -0.78 |
| Interpeduncular nucleus | IPN | Midbrain | C | 0 | 0.91 |
| Medial pretectal area | MPT | Midbrain | A | 0 | -0.15 |
| Midbrain reticular nucleus | MRN | Midbrain | B | 0.35 | 1.27 |
| Olivary pretectal nucleus | OP | Midbrain | A | 0.30 | -0.91 |
| Pedunculopontine nucleus | PPN | Midbrain | B | 0.45 | 0.43 |
| Periaqueductal gray | PAG | Midbrain | B | 0.50 | 0.12 |
| Precommissural nucleus | PRC | Midbrain | A | 0 | 0.17 |
| Substantia nigra, compact part | SNc | Midbrain | B | 0.14 | 0.78 |
| Substantia nigra, reticular part | SNr | Midbrain | B | 0.50 | 0.67 |
| Superior colliculus, motor related | SCm | Midbrain | B | 0.39 | 0.70 |
| Superior colliculus, sensory related | SCs | Midbrain | A | 0.38 | 0.90 |
| Ventral tegmental area | VTA | Midbrain | C | 0.03 | 0.57 |
| Pons | P | Hindbrain | B | 0.63 | -0.76 |
| Pons, motor related | P-mot | Hindbrain | B | 0.49 | -1.07 |
| Culmen | CUL | Cerebellum | A | 0 | -0.90 |

In each instance, the effect of group was driven by the 2BC/CIE group. Interestingly, the cortical amygdala/retrohippocampal and PSTN/TU regions were positively correlated with the amygdala group, whereas the other regions were negatively correlated.

### Clustering reveals a reorganization of co-activation networks during alcohol abstinence

We then performed data-driven hierarchical clustering of the distance matrices that were associated with the co-activation network of each condition to identify clusters of brain regions with similar patterns of co-activation (Lin et al., 2017, Wang et al., 2011, Kanonidis et al., 2016, Ray et al., 2017). Hierarchical clustering to 50% dendrogram height identified nine clusters for naive mice, eight clusters for 2BC/Air mice, and three clusters for 2BC/CIE mice (**Fig. 6)**. In the functional network for 2BC/CIE mice, an increase in coordinated activity was observed throughout the brain, reflected by an overall higher level of co-activation throughout the brain and larger clusters of regions that showed strong co-activation together.

**Figure 6.**
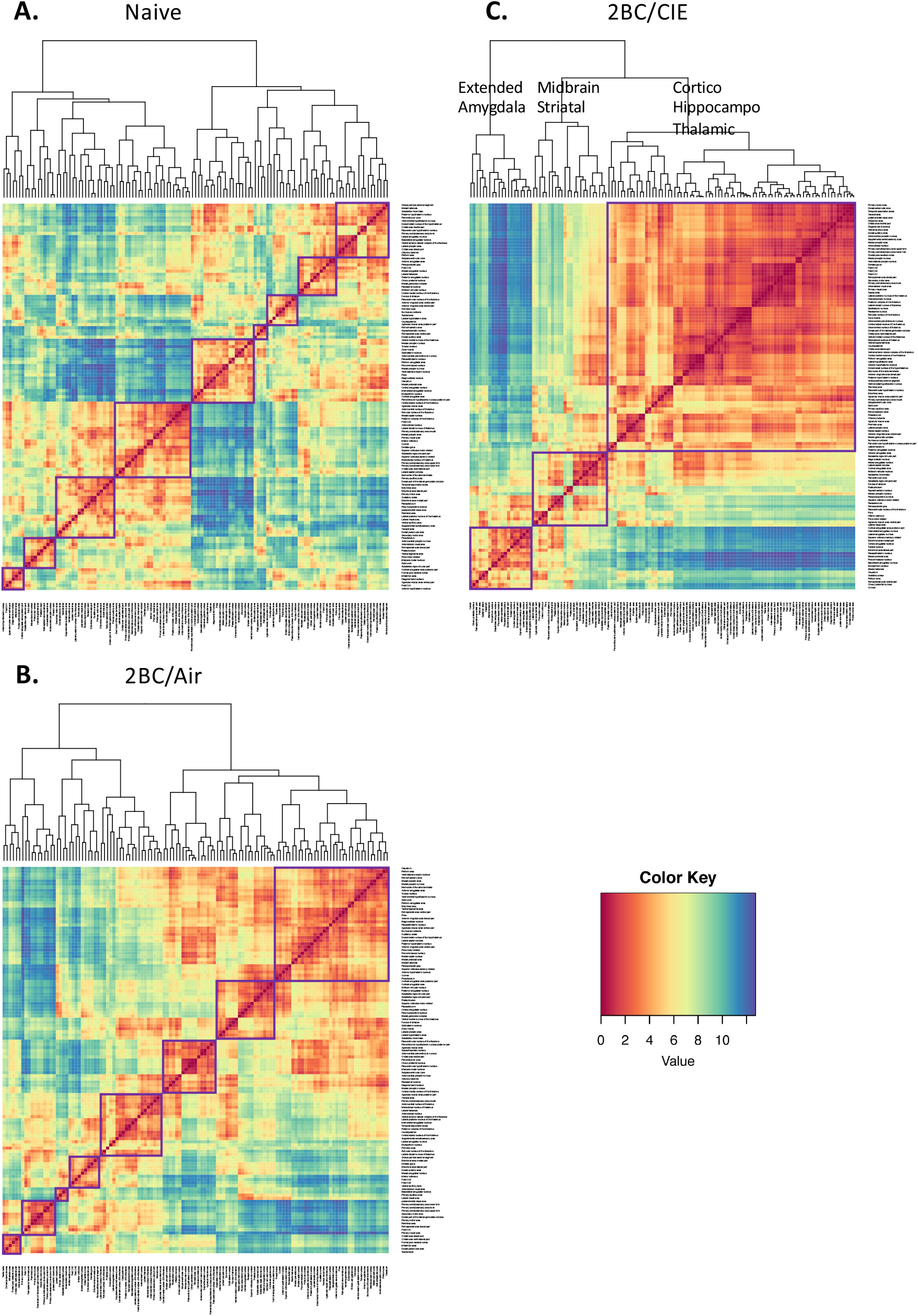
Hierarchical clustering of complete Euclidean distance matrices for each treatment. Clusters were determined by cutting each dendrogram at half of the maximal tree height. **A**. Relative distance of each brain region relative to the others examined in naive mice. In naive mice, we identified nine distinct clusters of co-activation. **B**. Relative distance of each brain region relative to the others examined in 2BC/Air mice. In 2BC/Air mice, we identified eight distinct clusters of co-activation. **C**. Relative distance of each brain region relative to the others examined in 2BC/CIE mice. In 2BC/CIE mice, we identified three distinct clusters of coactivation. For all distance matrices, each cluster is boxed in purple.

In the 2BC/CIE network, hierarchical clustering identified two clusters with opposing co-activation patterns and a third cluster that showed moderate co-activation with each of the other clusters. We named the clusters in the 2BC/CIE network based on the predominant regional components within each cluster (e.g., regions with the highest intracluster connectivity). One cluster was an extended amygdala cluster (cluster A), consisting of the CeA, BLA, LA, and IA. Additionally, this cluster contained the gustatory area (GU), PSTN, and medial habenula (MH), among other regions. A second cluster was a midbrain striatal cluster (cluster B), which consisted of the periaqueductal gray (PAG), paraventricular thalamus (PVT), pons, substantia nigra reticular and compact (SNr and SNc), MRN, and some additional amygdala areas that were not found in the extended amygdala cluster. The third cluster was a cortico-hippocampo-thalamic cluster (cluster C) that was highly anti-correlated with the extended amygdala cluster. The cortico-hippocampal-thalamic cluster included the PFC, OFC, sensory and somatosensory cortex, HIPP, THAL, and HYPO. This cluster also contained the VTA, IPN, lateral habenula (LH), bed nucleus of the stria terminalis (BST), and nucleus accumbens (ACB; **Fig. 6C**; see **Table 1** for a full list of regions and clusters).

### Identification of key brain regions associated with alcohol abstinence

To further characterize the functional network that is associated with alcohol abstinence, we identified key “hub” (e.g., the most inter- or intraconnected) brain regions of the network by graph theory analysis. We examined positive connectivity (thresholded to functional connections with a Pearson correlation coefficient of >0.75 [0.75R] for inclusion as a network connection) of the network that is associated with alcohol abstinence (2BC/CIE) using the clusters that were identified with hierarchical clustering to partition the regions of the network.

We examined individual brain regions in the network by looking at connections to other brain regions and determining the participation coefficient (PC; i.e., a measure of importance for intercluster connectivity) and the within-module degree Z-score (WMDz; i.e., a measure of importance for intracluster connectivity; (Guimera and Nunes Amaral, 2005). A high PC was considered ≥ 0.30, and a high WMDz was considered ≥ 0.70. Previous studies have used ranges of ≥ 0.30-0.80 for high PC and ≥ 1.5-2.5 for high WMDz (Guimera and Nunes Amaral, 2005, Cohen and D’Esposito, 2016). Because of differences in the sizes/types of networks that were examined, and methods used (e.g., Fos *vs*. functional magnetic resonance imaging [fMRI]) our WMDz values were considerably lower overall (ranging from −1.5 to 1.5). Therefore, we adjusted the range for high WMDz accordingly. This resulted in 7/20 regions for the extended amygdala cluster, 9/24 for the midbrain striatal cluster, and 25/79 for the hippocampal thalamic cluster that were considered to have high WMDz. Not all regions that fit these criteria are highlighted below, but a list of PC and WMDz values for all brain regions can be found in **Table 1**. We performed an overall assessment of the whole network and then focused on key regions from the extended amygdala cluster and its direct connections, given the importance of the extended amygdala in abstinence (Koob and Volkow, 2016, Koob and Le Moal, 1997, Koob and Volkow, 2010; **Fig. 7**).

**Figure 7.**
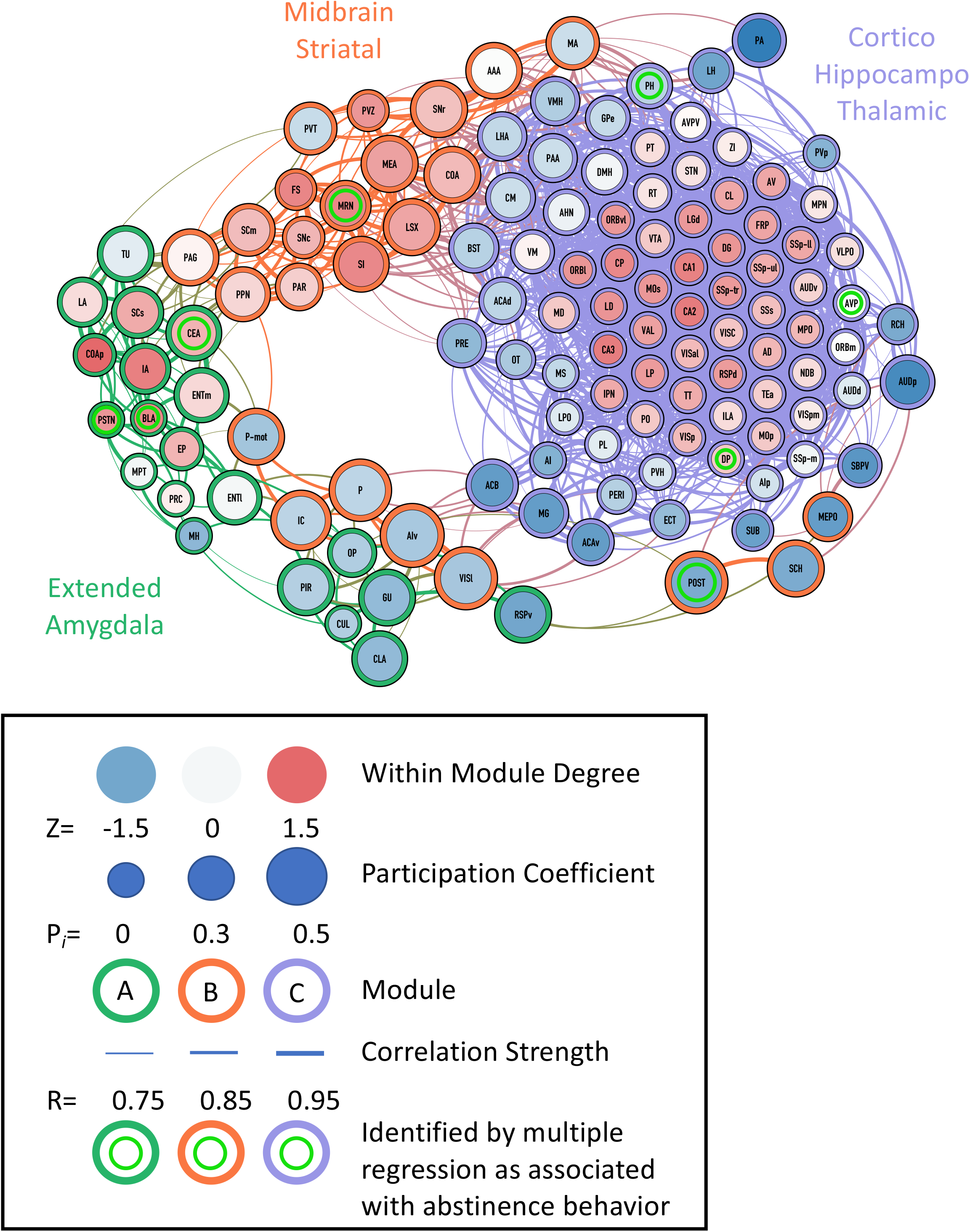
Functional connectivity of abstinence network thresholded to 0.75R. Nodes/brain regions of the network are represented by circles. The size of the node represents the participation coefficient (smaller = lower PC; larger = higher PC). The internal color of each circle represents the within-module degree Z-score (dark blue = lowest; dark red = highest). The color of the clusters identified in Fig. 6C are represented by different colored edges of each node circle (green = Cluster A; Red = Cluster B; Blue = Cluster C). The thickness of the lines represents the strength of the correlation between regions (thin = lower correlation; thick = higher correlation). Brain regions that are underlined with a color represent the regions that were identified by multiple regression (Fig. 3B) as important for the behaviors (yellow underline = drinking; purple underline = digging; teal underline = irritability-like behavior). See figure key for examples of each representative component of the figure.

We found that the extended amygdala cluster and cortico-hippocampo-thalamic cluster both interacted with the midbrain striatal cluster exclusively and not with each other. This was unsurprising because the extended amygdala cluster and cortico-hippocampo-thalamic cluster were found to have strong anti-correlations with each other. Interestingly, the cortico-hippocampo-thalamic cluster had numerous brain regions that drive coactivation within its own network, but none of these regions interacted with the other clusters. This is evident by the fact that the top 43 WMDz value brain regions of a total of 79 in the cluster had a PC of 0.10 or lower. Instead, a separate set of regions from the cortico-hippocampo-thalamic cluster was heavily involved in interactions with the midbrain striatal cluster, including such regions as the ACB, lateral hypothalamus (LHA), BST, and ventromedial hypothalamic nucleus (VMH), among others.

The midbrain striatal cluster had several regions that had both high PC and WMDz values, such as the medial amygdala (MEA), superior colliculus motor related (SCm), substantia innominata (SI), MRN, and others. Additionally, both the PAG and PVT had high PC values because of their exclusive connectivity with the extended amygdala cluster. The PAG especially acted as a strong hub between the extended amygdala and midbrain striatal clusters (high PC value connecting to regions in the extended amygdala cluster).

In the extended amygdala cluster, several brain regions had both high PC and WMDz values. These regions included the CEA, IA, and superior colliculus sensory related (SCs). Additionally, the COAp, PSTN, BLA, and EP had high WMDz values. Located near the major grouping of high WMDz regions of the extended amygdala cluster, the ENTm, LA, and TU had high PC values. Interestingly, several of these regions, including the TU, SCs, CeA, LA, and IA, connect strongly with the PAG, pedunculopontine nucleus (PPN), and PVT from the midbrain striatal cluster, and these regions collectively acted as the major bridge of connectivity between the two clusters.

Other regions in the extended amygdala cluster had high PC values connecting to the midbrain striatal cluster, such as the PIR, claustrum (CLA), GU, retrosplenial area ventral part (RSPv), and olivary pretectal nucleus (OP). These regions were connected to a separate group of regions from the midbrain striatal cluster that included the ventral agranular insula (AIv), lateral visual cortex (VISl), inferior colliculus (IC), P, POST, and suprachiasmatic nucleus (SCH).

## Discussion

The present study used single-cell whole-brain imaging and found that abstinence from alcohol dependence is associated with a reorganization of functional co-activation networks compared with nondependent and naive networks. Brain networks in dependent animals exhibited fewer larger clusters of co-activated brain regions, indicating that abstinence is associated with a reduction of modularity. This lower modularity resulted in the emergence of a new network architecture, in which the majority of brain regions were co-activated with each other in a cortico-hippocampo-thalamic cluster but anti-correlated with an extended amygdala cluster. Using graph theory, we identified candidate “hub” (i.e., high inter- or intracluster connectivity) regions of the extended amygdala cluster that may play a critical role in driving neural activity that is associated with alcohol abstinence. We complemented our functional network analysis with exploratory feature selection methods to identify a subset of brain regions that is most strongly predictive of addiction-like behaviors. These unbiased whole-brain analyses suggest that none of the mainstream theories of addiction alone is sufficient to explain neuronal network changes in addiction but instead provide evidence that favors the three-stage theory of addiction.

Using hierarchical clustering, we found a reduction of clustering in the 2BC/CIE functional network (three clusters) compared with the 2BC/Air (eight clusters) and naive (nine clusters) networks, indicating that alcohol abstinence results in a less modular brain overall. In humans, modularity has been found to decrease with age (Sporns and Betzel, 2016) and in cocaine addiction (Liang et al., 2015), and more modular networks are associated with better cognitive function (Gallen et al., 2016, Bertolero et al., 2018). Additionally, alcohol addiction is known to lead to cognitive dysfunction in humans and animal models (Golub et al., 2015, Parsons and Nixon, 1993, Sullivan et al., 2000, McKim et al., 2016, Le Berre et al., 2017, Crews and Boettiger, 2009). Changes in modularity that are caused by addiction are hypothesized to be long lasting and may help explain both cognitive dysfunction and relapse vulnerability.

We identified three modules that are associated with alcohol abstinence: (1) extended amygdala module, (2) midbrain striatal module, and (3) cortico-hippocampo-thalamic module. Notably, in some cases, brain regions that we would expect to be in one module were contained in another module, such as the BST, VTA, and ACB being part of the cortico-hippocampo-thalamic module. The extended amygdala and cortico-hippocampo-thalamic clusters had directly opposing activation patterns and included exclusively brain regions from specific anatomical groups. The brain regions in the extended amygdala cluster (e.g., CeA, IA, BLA, LA PSTN, TU, GU, MH, etc.), hypothesized to be involved in negative affect (Koob and Volkow, 2016), matched directly with the hedonic allostasis and three-stage theories of addiction. The cortico-hippocampo-thalamic cluster had opposing actions to the extended amygdala cluster and included almost all of the PFC, OFC, HIPP, sensory/somatosensory cortex, THAL, HYPO, VTA, and IPN. Several of these regions (e.g., PFC, OFC, insula, VTA, and ACB) match aspects from the habit, incentive salience, and three-stage theories of addiction. The midbrain striatal cluster contained several regions that are involved in midbrain-striatal dopamine reward signaling (e.g., SNr, SNc, PAG, MRN, lateral septal complex, SI, fundus of striatum, etc.), suggesting that this cluster fits well with aspects of the incentive salience (reward) and three-stage theories. Additionally, the ACB is one of the connector regions between the midbrain-striatal and cortico-hippocampo-thalamic clusters. The midbrain striatal cluster had moderate coactivation with both the extended amygdala and cortico-hippocampo-thalamic clusters, suggesting that this group of regions may act to integrate information from the other two clusters.

The extended amygdala has been heavily implicated in negative affect during drug withdrawal and protracted abstinence. However, much of the focus of previous studies has been on the CeA and BST (Pleil et al., 2016, Centanni et al., 2018, Erikson et al., 2018, Rinker et al., 2017, Pleil et al., 2015b, Koob, 2014, McBride, 2002, Smith and Aston-Jones, 2008, George et al., 2012b, Vendruscolo et al., 2012, Leao et al., 2015, de Guglielmo et al., 2016, Koob, 2008a, Koob, 2008b, Roberto et al., 2008, Francesconi et al., 2009, Roberto et al., 2010, Gilpin et al., 2011), and not on connections to other brain regions within this functional module. With regard to alcohol abstinence, the present findings suggest that as a functional module the extended amygdala may need to be redefined to include connections to additional brain areas, such as the PSTN, COAp, IA, EP, and BLA.

Using graph theory, we identified several non-CeA brain regions in the extended amygdala cluster that may drive alcohol abstinence. The TU was found to have high intercluster connectivity (high PC), primarily through the PAG, suggesting that the PAG and TU may communicate to drive neural activity during abstinence. The PAG has been shown to be involved in anxiety and hyperalgesia associated with alcohol abstinence (Avegno et al., 2018, Bonassoli et al., 2013), and chronic alcohol exposure has been found to alter PAG dopamine signaling (Li et al., 2013). The PSTN, COAp, EP, and BLA were found to have high intracluster connectivity (high WMDz), suggesting that they may drive activity within the extended amygdala circuit. Both the EP and BLA have been implicated in alcohol abstinence (Ruggeri et al., 2010, Morales et al., 2018, Diaz et al., 2011), but the PSTN and COAp remain understudied. The CeA, SCs, and IA presented both high intracluster and intercluster connectivity (high WMDz and PC), suggesting that these regions may be major drivers of abstinence symptoms and could be ideal targets for further study. The SC is involved in seizure responses during alcohol abstinence (Peris et al., 1992, Yang et al., 2001). The IA, although relatively understudied in alcohol addiction, contains dopamine D1 receptors (Fuxe et al., 2003), and an increase in D1 receptor density has been reported in alcohol-preferring rats following repeated alcohol deprivation (Sari et al., 2006).

Using stepwise multiple regression analysis, we identified the CeA as the main positive contributor to alcohol drinking and the MRN as a slightly negative contributor. As previously discussed, the CeA has been heavily implicated in alcohol abstinence and drinking. Identification of the CEA as a major contributor to alcohol drinking behavior validates our approach. The MRN is involved in sleep, wakefulness, and attention (Brown et al., 2012), which may indirectly moderate alcohol drinking. We did not identify regions other than the CeA that were previously implicated in alcohol drinking (Koob and Volkow, 2016, Hwa et al., 2017), however this is likely attributable to the multiple regression approach that uses stringent stepwise elimination of brain regions with redundant predictive values. Additionally, differences in behavioral paradigms may also explain why some regions that have been previously implicated in excessive alcohol drinking (e.g., BST Lovinger and Kash, 2015)] or MH (Kononoff et al., 2018)) were not identified in the present study via multiple regression.

We identified the PH, PSTN, and DP as the main positive contributors to irritability-like behavior. The blockade of corticotropin-releasing factor-1 (CRF1) receptors has been shown to reduce irritability-like behavior in rats (Kimbrough et al., 2017). The PH, PSTN, and DP all express CRF or CRF receptors (Gao et al., 2016, Zhu et al., 2012, Ketchesin et al., 2017, George et al., 2012b). Additionally, activation of the DP during alcohol abstinence predicts cognitive impairment and excessive alcohol drinking (George et al., 2012b). Thus, the PH, PSTN, and DP are potential targets for the future exploration of irritability-like behavior that is caused by alcohol abstinence.

We identified the BLA, AVP, and POST as the main positive contributors to digging behavior. Numerous studies of alcohol drinking and abstinence have used digging behavior as a measure of affective dysfunction (Sidhu et al., 2018, Jury et al., 2017, Pleil et al., 2015a, Rose et al., 2016, Umathe et al., 2008), and the BLA has been correlated with digging behavior (Palazzo et al., 2015). Overall, however, the BLA, AVP, and POST may be relatively novel targets to explore with regard to their contribution to digging behavior, especially affective dysfunction during alcohol abstinence.

Finally, three of the brain regions that were identified by stepwise multiple regression as relevant to abstinence-related behaviors (i.e., increases in drinking, irritability, and digging) are part of the extended amygdala cluster that was identified by hierarchical clustering (CeA, BLA, and PSTN), further emphasizing the importance of this group of brain regions.

A known limitation of the current study is the focus on the neuronal networks of abstinence. It is likely that the network structure identified here would be different at other time points (intoxication, acute withdrawal, relapse). Follow up studies will be critical to further understand, not only the structure of the neuronal network of abstinence, but also its dynamic during the different phases of the addiction cycle.

The present study demonstrates that abstinence from alcohol dependence is associated with a reduction of modularity of the brain into three major functional modules, two with opposing co-activation patterns. This lower modularity resulted in the emergence of a new network architecture that is composed of three major modules: cortico-hippocampo-thalamic module that was anti-correlated with an extended amygdala module and both modules that were connected via an intermediate midbrain striatal module. The present data match the three-stage theory of addiction (Koob and Volkow, 2016, Koob and Le Moal, 1997, Koob and Volkow, 2010) better than any single theory (incentive salience, hedonic allostasis, habit). Within one of the clusters identified (i.e., an extended amygdala cluster), we found brain regions that are associated with negative affect, the activation of which predicted alcohol abstinence-dependent behaviors. Among the regions that were identified in the extended amygdala cluster, the PSTN, TU and IA were identified as potential drivers of the network and represent novel targets for future studies of alcohol abstinence. Altogether, these results provide a unique brain map of alcohol dependence, identify lower network modularity as a key feature of alcohol dependence, and identify three main modules that drive addiction-like behaviors in accordance with the three-stage hypothesis of addiction.

## Materials and Methods

### Animals

Age-matched male C57BL/6J mice that were bred at The Scripps Research Institute (20-30 g) were used for the experiments. The mice were single-housed for the entire duration of the study. The mice were maintained on a 12 h/12 h light/dark cycle with *ad libitum* access to food and water with 7090 Teklad sani-chips (Envigo) as bedding for the home cages and experimentation. All of the procedures were conducted in strict adherence to the National Institutes of Health Guide for the Care and Use of Laboratory Animals and approved by The Scripps Research Institute Institutional Animal Care and Use Committee.

### Behavioral testing

#### Two-bottle choice/chronic intermittent ethanol vapor

We used the 2BC/CIE paradigm, a well-established mouse model of alcohol dependence (Contet et al., 2011, Kreifeldt et al., 2013, Becker and Lopez, 2004, Gorini et al., 2013), to induce the escalation of alcohol drinking and behavioral symptoms of abstinence (see **Fig. 1B**). In the 2BC/CIE paradigm, weeks of voluntary ethanol drinking during limited-access 2BC sessions are alternated with weeks of CIE.

During 2BC weeks, the mice were given access to two bottles that contained water and alcohol (15%, v/v), respectively, Monday-Friday, for 2 h, starting at the beginning of the dark phase of the circadian cycle. During CIE weeks, the mice were exposed to four cycles of 16-h intoxication/8-h abstinence (Monday-Friday) followed by 72-h abstinence (Friday-Monday). Each 16-h period of alcohol vapor exposure was primed with an intraperitoneal injection of alcohol (1.5 g/kg) to initiate intoxication and pyrazole (an alcohol dehydrogenase inhibitor, 1 mmol/kg) to normalize alcohol clearance rate between individual mice. Average blood alcohol levels were measured periodically at the end of alcohol vapor inhalation periods and averaged (164.6 ± 17.0 mg/dl). Air-exposed mice received injections of pyrazole only.

The mice were first given 2 weeks of 2BC (weeks 1-2) and were then split into two groups of equivalent baseline intake (2BC/Air, *n* = 5; 2BC/CIE, *n* = 4). The mice were then subjected to five rounds of alternating weeks of Air/CIE exposure with weeks of 2BC drinking (weeks 3-12), followed by an additional 6^th^ week of Air/CIE exposure (week 13). The mice were tested for irritability and digging behaviors 7 and 10 days, respectively, into abstinence from vapor (week 14) and were then exposed to a final 7^th^ week of Air/CIE inhalation (week 15). They were perfused 7 days after the last vapor exposure, with no intervening behavioral testing or voluntary drinking, at the same time of day as 2BC sessions, for brain collection (week 16). Brains from age-matched single-housed alcohol-naive mice (*n* = 5) were also collected at the same time.

### Irritability-like behavior

Irritability-like behavior was assessed using the bottle brush test (BBT), conducted as described by Riittinen et al. (Riittinen et al., 1986). The BBT measures defensive and aggressive responses to an “attack” by a mechanical stimulus (a moving bottle brush, Lagerspetz and Portin, 1968). The BBT has recently been used in alcohol-dependent rats and mice to identify increases in irritability-like behavior during alcohol abstinence (Somkuwar et al., 2017, Kimbrough et al., 2017, Sidhu et al., 2018). The methods were similar to Sidhu et al. (2018). Testing was conducted under red light, and the mouse was “attacked” by moving a bottle brush (14 cm length × 5 cm width cylindrical brush, 33 cm total length with handle) toward it. The attacks were made in the home cage with the lid and food tray removed. Each test consisted of 10 trials with 10 s intertrial intervals. Briefly, the mouse started each trial at the opposite end of the cage and was then “attacked” by the brush. Each attack consisted of five stages: (1) the brush rotating toward the mouse from the opposite end of the cage, (2) the brush rotating against the whiskers of the mouse, (3) the brush rotating backward toward the starting position in the opposite end of the cage, (4) the brush rotating at the starting position, and (5) the brush at the starting position not rotating. Each stage lasted 1.5 s, with the exception of stage 5 that was prolonged, if necessary, until the mouse returned to its end of the cage or 5 s had elapsed. Responses to the attacks were observed. The following behavioral responses were scored: smelling the brush, exploring the brush, biting the brush, boxing the brush, following the brush, tail rattling, escaping from the brush, digging, jumping, climbing/rearing, defecation, vocalization, and grooming. The total number of occurrences of each behavior across all 10 trials was recorded and summed to calculate a total irritability-like behavior score.

### Digging

Digging and marble burying (DMB) were measured as described by Deacon and Sidhu et al. (Deacon, 2006, Sidhu et al., 2018). Testing was conducted under dim lighting (20 lux). The mouse was placed in a new, clean cage with a bedding thickness of 5 cm and no lid and allowed to freely dig for 3 min. The number of digging bouts and total digging duration were recorded (phase 1). The mouse was then removed from the cage. The bedding was flattened, and 12 marbles were arranged in a 4 × 3 array on top of the bedding. The mouse was reintroduced into the cage and allowed to bury the marbles for 30 min with a lid that covered the cage. The number of marbles that were buried (covered two-thirds or more by bedding) was counted at the end of the test (phase 2).

### Tissue collection

The mice were deeply anesthetized and perfused with 15 ml of phosphate-buffered saline (PBS) followed by 50 ml of 4% formaldehyde. The brains were postfixed in formaldehyde overnight. The next day, brains were washed for 30 min three times with PBS and transferred to a PBS/0.1% azide solution at 4°C for 2-3 days before processing via iDISCO.

### iDISCO

The iDISCO procedure was performed as reported by Renier et al. (Renier et al., 2016, Renier et al., 2014).

### Immunostaining

Fixed samples were washed in 20% methanol (in double-distilled H2O) for 1 h, 40% methanol for 1 h, 60% methanol for 1 h, 80% methanol for 1 h, and 100% methanol for 1h twice. The samples were then precleared with overnight incubation in 33% methanol/66% dichloromethane (DCM; Sigma, catalog no. 270997-12X100ML). The next day, the samples were bleached with 5% H_2_O_2_ (1 volume of 30% H_2_O_2_ for 5 volumes of methanol, ice cold) at 4°C overnight. After bleaching, the samples were slowly re-equilibrated at room temperature and rehydrated in 80% methanol in (double-distilled H_2_O) for 1 h, 60% methanol for 1 h, 40% methanol for 1 h, 20% methanol for 1 h, PBS for 1 h, and PBS/0.2% TritonX-100 for 1 h twice. The samples were then incubated in PBS/0.2% TritonX-100/20% dimethylsulfoxide (DMSO)/0.3 M glycine at 37°C for 2 days and then blocked in PBS/0.2% TritonX-100/10% DMSO/6% donkey serum at 37°C for 2 days. The samples were then incubated in rabbit anti c*fos* (1:500; Santa Cruz Biotechnology, catalog no. sc-52) in PBS-0.2% Tween with 10 μg/ml heparin (PTwH)/5% DMSO/3% donkey serum at 37°C for 7 days. The samples were then washed in PTwH for 24 h (five changes of the PTwH solution over that time) and incubated in donkey anti-rabbit-Alexa647 (1:500; Invitrogen, catalog no. A31573) in PTwH/3% donkey serum at 37°C for 7 days. The samples were finally washed in PTwH for 1 day before clearing and imaging.

### Sample clearing

Immunolabeled brains were cleared using the procedure of Reiner et al. (2016). The samples were dehydrated in 20% methanol (in double-distilled H_2_O) for 1 h, 40% methanol for 1 h, 60% methanol for 1 h, 80% methanol for 1 h, 100% methanol for 1 h, and 100% methanol again overnight. The next day, the samples were incubated for 3 h in 33% methanol/66% DCM until they sank to the bottom of the incubation tube. The methanol was then washed for 20 min twice in 100% DCM. Finally, the samples were incubated in DiBenzyl Ether (DBE; Sigma, catalog no. 108014-1KG) until clear and then stored in DBE at room temperature until imaged.

### Image acquisition

Left hemispheres of cleared samples were imaged in the sagittal orientation (right lateral side up) on a light-sheet microscope (Ultramicroscope II, LaVision Biotec) equipped with a sCMOS camera (Andor Neo) and 2×/0.5 objective lens (MVPLAPO 2×) equipped with a 6 mm working distance dipping cap. Imspector Microscope controller v144 software was used. The microscope was equipped with an NKT Photonics SuperK EXTREME EXW-12 white light laser with three fixed light sheet generating lenses on each side. Scans were made at 0.8× magnification (1.6× effective magnification) with a light sheet numerical aperture of 0.148. Excitation filters of 480/30, 560/40, and 630/30 were used. Emission filters of 525/50, 595/40, and 680/30 were used. The samples were scanned with a step-size of 3 μm using dynamic horizontal scanning from one side (the right) for the 560 and 630 nm channels (20 acquisitions per plane with 240 ms exposure, combined into one image using the horizontal adaptive algorithm), and without horizontal scanning for the 480 nm channel using two-sided illumination (100 ms exposure for each side, combined into one image using the blending algorithm). To speed up acquisition, both channels where acquired in two separate scans. To account for micro-movements of the samples that may occur between scans, three-dimensional image affine registration was performed to align both channels using ClearMap (Renier et al., 2016). Images for coronal representative figures were captured with similar settings, except the brains were oriented with the olfactory bulbs up during acquisition, and acquisition used both left and right light sheets with horizontal scanning and a blending algorithm.

### Data analysis

Behavioral data were analyzed, and Pearson correlations were calculated using Statistica software (Tibco). Hierarchical clustering was performed using R Studio software.

### Behavioral assessments

Drinking data are presented as average weekly intake for 2BC sessions. The drinking data were analyzed using repeated-measures ANOVA of the average baseline intake and the last two weeks of drinking. *Post hoc* analysis was performed using the Student-Newman-Keuls test. 2BC/Air mice and naive mice did not show significant differences in any measure of irritability-like behavior or digging behavior and thus were combined into a single control group for analyses of these data. Irritability-like behavior and digging behavior were analyzed using *t*-tests. Values of *p* < 0.05 were considered significant for 2BC/CIE *vs*. control mice. Digging behavior (number of bouts, duration of bouts, and number of marbles buried) was combined into a single digging value by calculating the Z-score for each individual behavior for all animals across all treatments and then obtaining the average Z-score across all three behaviors for each animal.

### Identification of activated brain regions

Images that were acquired from the light-sheet microscope were analyzed from the end of the olfactory bulbs (olfactory bulbs not included in the analysis) to the beginning of the hindbrain and cerebellum. Counts of Fos-positive nuclei from each sample were identified for each brain region using ClearMap (Renier et al., 2016). ClearMap uses autofluorescence that is acquired in the 488 channel to align the brain to the Allen Mouse Brain Atlas (2004) and then registers Fos counts to regions that are annotated by the atlas. The data were normalized to a log10 value to reduce variability and bring brain regions with high numbers (e.g., thousands) and lower numbers (e.g. tens to hundreds) of Fos counts to a similar scale.

### Behavioral comparisons to neural activation

Forward stepwise multiple regressions were performed for each behavior/log10 Fos combination to determine which brain regions may account for the majority of variability for the given behavior. We first used very inclusive values for the model (data not shown) and then adjusted the tolerance and *F* to enter values to only include brain regions that represented ≥ 10% of the variability. The final forward step-wise analysis was performed using a 0.5 tolerance value and a *F*-to-enter-value of 6.7. The *F*-to-enter-value corresponded to *t* = 2.6.

### Identification of co-activation within individual networks

Separate inter-regional Pearson correlations were then calculated across animals for the 2BC/CIE, 2BC/Air, and naive groups to compare the log10 Fos data from each brain region to each of the other brain regions. For all of the functional co-activation network analyses, the 2BC/Air and naive groups were analyzed separately to maintain a relatively equal *n* (*n* = 4 for 2BC/CIE, *n* = 5 for 2BC/Air, and *n* = 5 for naive) for the correlation calculations. Instead of using an alphabetical arrangement of each anatomical group from the Allen Mouse Brain Atlas (2004), we split 2BC/CIE inter-regional Fos correlations into individual anatomical groups (i.e., cortical plate, cortical subplate, striatum, pallidum, thalamus, hypothalamus, and midbrain plus hindbrain). We then calculated the complete Euclidean distance and performed hierarchical clustering of each individual Allen Mouse Brain Atlas group separately. We then rearranged the order of the brain regions for each group based on hierarchical clustering. Using this order for each anatomical Allen Mouse Brain Atlas group, we then merged the groups back together, resulting in an “ordered Allen Mouse Brain Atlas list,” which was then applied to arrange the heatmaps of correlations for all treatments (**Fig. 5A-C**). This arrangement did not alter the values in any way and was used solely for visualization purposes.

### Analysis of amygdala cluster vs. other major brain clusters

For each treatment condition, average *R* values were calculated for correlations between each individual brain region of the amygdala cluster and all brain regions from each of the other clusters that were examined (e.g., average *R* for CEA to cortical regions). An average and standard error of the mean were then calculated across the average *R* values for all of the amygdala brain regions for each given comparison. A one-way ANOVA was then performed to examine the effect of treatment condition on the average *R* value for each amygdala *vs*. other cluster comparisons (e.g., average *R* for amygdala to cortical regions).

### Hierarchical clustering

Inter-regional Fos correlations were then used to calculate complete Euclidean distances between each pair of brain regions in each group of mice. The distance matrices were then hierarchically clustered by both row and column using the complete method to identify clusters of co-activation within each treatment group. The hierarchical cluster dendrograms were trimmed at half the height of each given tree to split the dendrogram into specific clusters.

### Graph theory identification of functional networks

We used a graph theory-based approach to identify the functional neural network of abstinence symptoms that are seen in alcohol dependence. Graph theory is a branch of mathematics that is used to analyze complex networks, such as social, financial, protein, and neural networks (Jeong et al., 2001, Barabasi, 2009, Babu et al., 2012, Vetere et al., 2017, Wheeler et al., 2013, Jarrell et al., 2012, Oh et al., 2014, Varshney et al., 2011, Markov et al., 2014, Chiang et al., 2011, Bullmore and Sporns, 2012, Bargmann and Marder, 2013, Cohen and D’Esposito, 2016). Using graph theory, functional networks can be delineated, and key brain regions of the network can be identified (Rubinov and Sporns, 2010, Sporns et al., 2007, Wheeler et al., 2013, Vetere et al., 2017).

Previous studies of regional connectivity profiles in Fos co-activation networks focused on global measures of connectivity (e.g., degree, Wheeler et al., 2013). However, in correlation-based networks, these measures can be strongly influenced by the size of the sub-network (cluster) in which a node participates (Power et al., 2013). For the graph theory analyses, we were interested in regional properties and not cluster size *per se*. Thus, it is necessary to consider cluster structure when examining the role that each region plays in the network. To accomplish this, we utilized two widely used centrality metrics that were designed for application to modular systems. The WMDz indexes the relative importance of a region within its own cluster (e.g., intracluster connectivity), and the PC indexes the extent to which a region connects diversely to multiple clusters (e.g., intercluster connectivity; (Guimera and Nunes Amaral, 2005).

We first took the Pearson correlation values that were calculated for the brain regions from 2BC/CIE mice. Prior to plotting and calculating regional connectivity metrics, the network was thresholded to remove any edges that were weaker than *R* = 0.75. As such, visualization and graph theory analyses were performed using only edges with positive weights. Regional connectivity metrics (PC and WMDz) were calculated as originally defined by Guimerà and Amaral (2005), modified for application to networks with weighted edges. PC and WMDz were calculated using a customized version of the bctpy Python package (https://github.com/aestrivex/bctpy), which is derived from the MATLAB implementation of Brain Connectivity Toolbox (Rubinov and Sporns, 2010).

For WMDz, let *k_i_* (within-module degree) be the summed weight of all edges between region *i* and other regions in cluster *s_i_*. Then, *k̅_s_i__* is the average within-module degree of all regions in cluster *s_i_*, and *σ_k_s_i___* is the standard deviation of those values. The Z-scored version of within-module degree (WMDz) is then defined as:

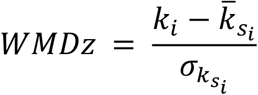

This provides a measure of the extent to which each region is connected to other regions in the same cluster.

For PC, let *k_is_* (between-module degree) be the summed weight of all edges between region *i* and regions in cluster *s*, and *k_i_* (total degree) be the summed weight of all edges between region *i* and all other regions in the network. The PC of each region is then defined as:

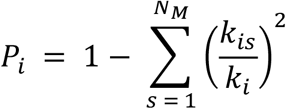

This provides a measure of the extent to which the connections of a region are distributed mostly within its own cluster (PC approaching 0) or distributed evenly among all clusters (PC approaching 1).

Network visualization was performed using a combination of Gephi 0.9.2 software (Bastian et al., 2009) and Adobe Illustrator software. Nodes were positioned using the Force Atlas 2 algorithm (Jacomy et al., 2014).

## Supporting information

## Acknowledgements

The authors thank Dr. Nicolas Renier for technical guidance, Michael Arends for editorial assistance, and Lauren C. Smith for assistance with illustrations. Light sheet imaging was performed at the Cal-Tech Beckman Institute.

## CREdit Taxonomy Author Contributions

Conceptualization: A.K. and O.G. Methodology: A.K., C.C., and O.G. Formal Analysis: A.K., D.J.L., M.D., C.C., and O.G. Investigation: A.K., A.C., H.S., and M.K. Resources: O.G., A.C., and C.C. Writing-Original Draft: A.K. and O.G. Writing-Review & Editing: A.K., A.C., D.J.L., M.D., C.C., and O.G. Visualization: A.K., A.C., and D.J.L. Supervision: M.D. Funding Acquisition: A.K., O.G., A.C., M.D., and C.C.

