## Supplementary figures and images for "Single-cell whole-brain imaging and network analysis provide evidence of the three-stage hypothesis of addiction"

### Supplementary file 1

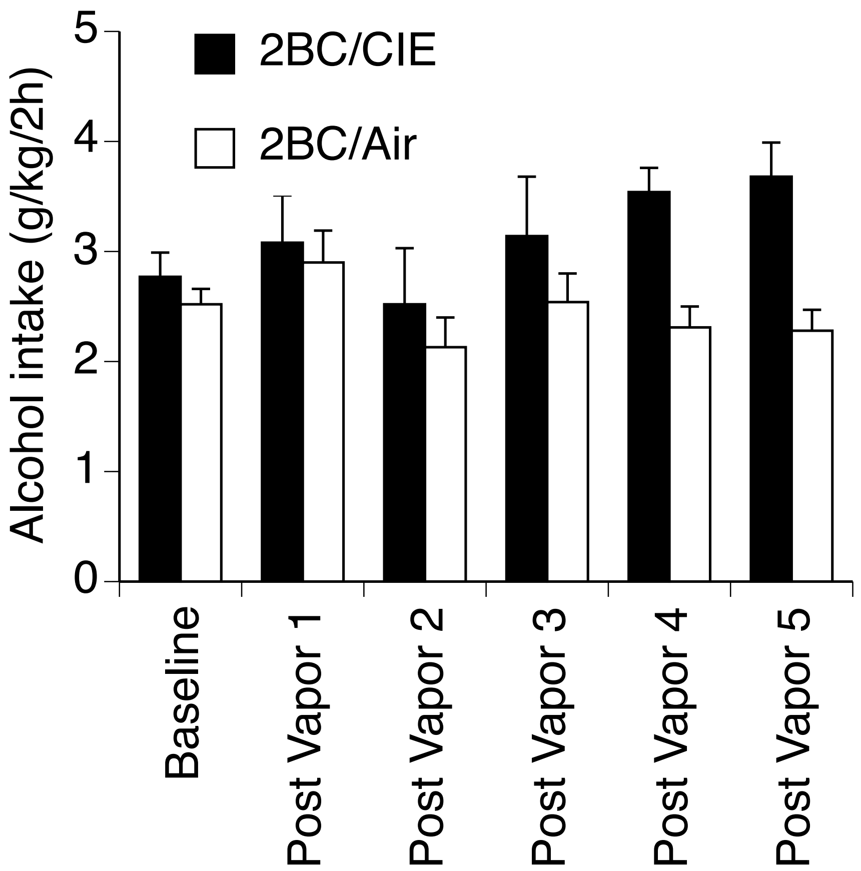


Alcohol drinking in 2BC/CIE (black bars) and 2BC/Air (white bars) mice across all weeks of drinking.
