## Supplementary material for "Single-cell whole-brain imaging and network analysis provide evidence of the three-stage hypothesis of addiction"

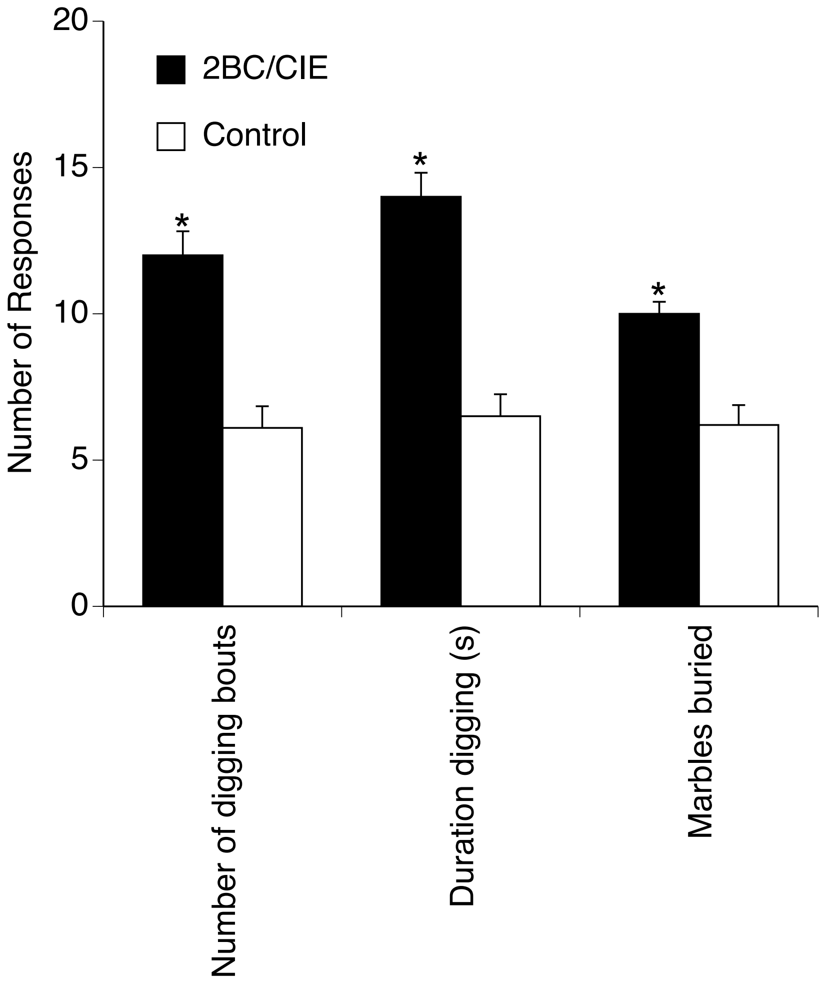


Individual digging behaviors for 2BC/CIE (black bars) and control (white bars) mice. 2BC/CIE mice had increased number of digging bouts, duration digging and marbles buried.
