## Supplementary material for "Single-cell whole-brain imaging and network analysis provide evidence of the three-stage hypothesis of addiction"

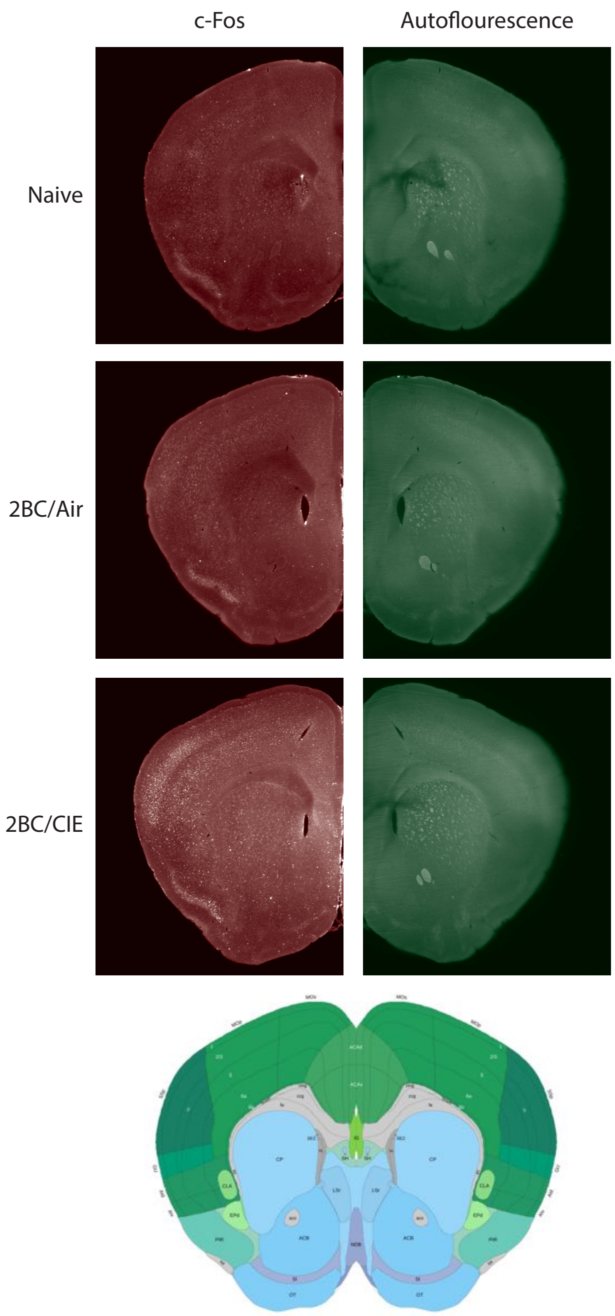


Example coronal image from each treatment showing c-Fos expression and autofluorescence used to automatically register and align the images to the Allen Brain Atlas.

Supplemental Figure 3
